# Enformer-Based Phylogenetic Tree Reconstruction

**DOI:** 10.64898/2026.06.05.730316

**Authors:** Alessandra Capitanio, Nicolò Stranieri, Alessio Enderti, Simone G. Riva, Jim R. Hughes

**Author notes:** These authors contributed equally to this work (Alessandra Capitanio, Nicolò Stranieri).

## Abstract

Enformer is a deep learning model trained on human and mouse genomes to predict regulatory activity from 196,608 bp DNA windows. Its trunk embeddings capture long-range cis-regulatory interactions, but whether this signal generalises across the tree of life has not been assessed.

We embed universal single-copy orthologous groups (OGs) from OrthoDB v12 across three taxonomic scales and evaluate reconstructed trees against TimeTree5 using Mantel *r* and Normalised Robinson-Foulds (NRF). On 702 OGs across 34 Primate species (≤ 74 Mya), the consensus tree achieves Mantel *r* = 0.902 and NRF = 0.481, correctly recovering major clades. A key finding is that flanking regulatory context — not the gene locus itself — carries the phylogenetic signal: restricting pooling to central 448 bins collapses Mantel *r* to 0.355. Applying the same fixed configuration to Vertebrates (≤ 450 Mya, 83 OGs, 150 species) and Plants (≤ 1,500 Mya, 92 OGs, 40 species) yields consensus Mantel *r* of 0.752 and 0.803 respectively, with NRF worsening monotonically across tiers.

Distance-ordering fidelity degrades smoothly with evolutionary distance while topological accuracy declines steadily, with no sharp taxonomic boundary. These results show that an unmodified regulatory deep learning model encodes robust phylogenetic signal well beyond its training distribution, reaching across 1,500 million years of divergence.

## I. Introduction & Motivation

Phylogenetics is the study of evolutionary relationships among biological entities, including species, individuals, or genes. By leveraging DNA and RNA sequence data, it seeks to reconstruct evolutionary history, which is typically represented as branching diagrams known as phylogenetic trees (or phylogenies). Phylogenetic reconstruction methods aim to infer these relationships by modelling sequence evolution. Traditionally, phylogenetic reconstructions have relied on multiple sequence alignment (MSA) techniques, taking as a basis the orthologous marker genes, and assessing the divergence between species by distance-based, maximum-likelihood, or Bayesian inference [1]. MSA requires direct comparison between nucleotide sequences and assumes positional homology, an assumption often violated by genome rearrangement or extreme divergence between species.

Beyond alignment-based strategies, several alternative approaches have been proposed to overcome these limitations, particularly k-mer–based methods. These approaches represent sequences as vectors of substring (k-mer) frequencies and estimate similarity without requiring positional homology, offering improved scalability and robustness for highly divergent or rearranged genomes. However, k-mer methods inherently discard positional and contextual information, and genome-wide biases such as GC content or the presence of repetitive elements can introduce spurious similarities that do not reflect shared ancestry. In addition, they are unable to capture long-range dependencies and higher-order regulatory structure within biological sequences [2].

Recent advances in deep learning seek to address these limitations by learning hierarchical, context-aware representations directly from raw DNA sequences. These methods aim to capture complex, non-linear patterns that are not accessible to k-mer statistics or alignment-based models. For instance, DEPP [3] learns a neural embedding of sequences in which Euclidean distances approximate phylogenetic distances, enabling placement of new sequences onto a reference tree. DeePhy [4] applies neural networks to infer small subtree structures, such as triplets, as part of phylogenetic reconstruction. In addition, methods such as Quartformer [5] reformulate phylogenetic inference as a supervised learning problem over small substructures, predicting quartet topologies using transformer-based architectures and assembling these local predictions into a global tree.

These approaches primarily operate directly on raw sequence similarity and evolutionary signal, without explicitly incorporating higher-order biological context encoded in functional genomics. In contrast, recent foundation models such as Enformer have demonstrated the ability to learn rich representations of DNA sequences that capture long-range regulatory interactions and functional genomic constraints across kilobase-scale regions [6].

Enformer is a deep learning model trained on human and mouse genomes. It takes DNA sequences of 196,608 base pairs as input and processes them through a convolutional–transformer architecture to predict gene expression and regulatory activity. The model captures genomic context over distances of up to approximately 100 kb, enabling it to integrate both local motifs and long-range regulatory interactions. Its intermediate “trunk” representation (of shape 896 × 3072, before the prediction heads) encodes rich sequence features that reflect interactions between cis-regulatory elements. Because these embeddings are learned from functional genomic signals, they may also implicitly capture sequence constraints shaped by evolutionary divergence across species [6].

In this work, we investigate whether Enformer can be leveraged to reconstruct phylogenetic trees by embedding orthologous sequences and assess its ability to generalise across genome-scale data spanning a wide range of evolutionary distances. We do not benchmark against existing phylogenetic methods; our goal is not to outperform them, but rather to test whether a regulatory foundation model encodes phylogenetic signal at all, and to characterise Enformer’s representational capacity as a genomic foundation model.

## II. Methods

### A. Pipeline

For every taxonomic level, we apply the same six-step pipeline: (i) retrieve universal single-copy OGs from OrthoDB [7] at the relevant taxonomic node; (ii) resolve genomic coordinates from NCBI [8] for each (OG, species) pair; (iii) construct a symmetric 196,608 bp window centred on each gene; (iv) extract Enformer trunk embeddings; (v) pool the per-(OG, species) trunk output into a single embedding vector; and (vi) evaluate phylogenetic reconstruction quality against TimeTree5 [9]. The pooling strategy used in step (v) is the one identified as optimal on the Primate benchmark and is held fixed at all broader levels. Only the OrthoDB taxon identifier differs between runs.

#### (i) Orthologous Group Selection

OGs are queried from OrthoDB v12 through its REST API with filters `universal=1` (present in every species at the queried node) and `singlecopy=1` (single-copy in every species). When it is not possible for all species at that level, we will select a subset of OGs and species to satisfy this condition. Node identifiers follow NCBI taxonomy: Primates (9443), Vertebrata (7742) and Viridiplantae (33090).

#### (ii) Gene Coordinate Retrieval

Each OrthoDB entry carries a `pub_gene_id` field in one of three forms, each resolved through a distinct NCBI endpoint: numeric Gene IDs are submitted to `esummary` in batches of 500; gene symbols are resolved through `esearch` then `esummary`; and genome-assembly accessions are queried via the NCBI Datasets v2 API in batches of 50 symbols per taxon. Species lacking annotation in NCBI are excluded. Coordinates are normalised by taking the min/max of strand endpoints, and genes whose genomic span exceeds the Enformer output-region width (114,688bp) are discarded. Only OGs represented in every retained species are kept, or a subset of species is selected to satisfy this condition.

#### (iii) Enformer Input Window Construction

For each retained (OG, species) entry, the input window is centred on the gene mid-point:

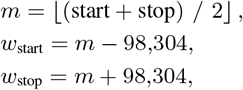

yielding a symmetric 196,608 bp window, matching the Enformer input context used for regulatory prediction [6]. Windows that exceed a chromosome boundary are padded with `N` characters on the affected side. DNA sequences are fetched from NCBI `nuccore` via `efetch`.

#### (iv) Enformer Trunk Embedding Extraction

Sequences are one-hot encoded (A → [1, 0, 0, 0], C → [0, 1, 0, 0], G → [0, 0, 1, 0], T→ [0, 0, 0, 1], N → [0, 0, 0, 0]) and forwarded through Enformer with gradient tracking disabled. A forward hook on the final trunk layer (immediately before the human/mouse prediction heads) captures the intermediate representation of shape (*B ×* 896 *×* 3072), where *B* is the number of species in the OG processed as a single GPU mini-batch [6]. All experiments were run on a single NVIDIA L40S GPU with a mini-batch size of *B* = 16. Raw trunk tensors are saved as compressed `.npz` files so that all downstream strategy comparisons reuse the same cached activations.

#### (v) Embedding Strategies

Five pooling strategies collapse the 896 *×* 3072 trunk output into a single *d*-dimensional vector per (OG, species) entry:

- **A — mean all bins:** 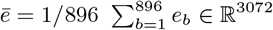
- **B — mean central 448 bins:** bins 224–671 (0-indexed, half-open), the central half of the window.
- **C — mean gene bins:** bins overlapping the annotated gene locus (≈ 10–50 bins, gene-length dependent).
- **D — mean + std:** [ē, *σ*_*e*_] ∈ ℝ^6144^.
- **E — max all bins:** *e*_*j*_ = max_*b*_(*e*_*b,j*_).

For strategy A we additionally evaluate PCA reduction [10] to *n*_comp_ ∈ {16, 32} components in two modes. In *per-OG PCA*, we fit a separate PCA on each OG’s *n*_species_ *×* 3072 matrix after `StandardScaler` z-score normalisation. In *global PCA*, a single PCA is fit once on all OGs stacked into an (*n*_OG_ · *n*_species_) *×* 3072 matrix and the same projection is applied to every OG. Each configuration is evaluated under three distance metrics: Euclidean, cosine, and correlation.

The strategy search is conducted exclusively on the Primate benchmark. The winning configuration is then held fixed and applied unchanged to all broader taxonomic levels.

### B. Evaluation Metrics

#### Mantel r

It is the Pearson correlation [11] between the upper triangle of the pairwise embedding-distance matrix and the corresponding pairwise TimeTree5 divergence-time matrix [12]. Pairs lacking a TimeTree5 entry are excluded. Higher values indicate stronger agreement with the molecular clock.

#### Normalised Robinson-Foulds (NRF)

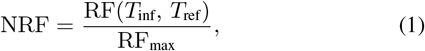

where RF is the symmetric difference of the bipartition sets of the inferred tree *T*_inf_ and the reference tree *T*_ref_ [13], and RF_max_ is returned by `ete3` for the corresponding leaf set. *T*_inf_ is an average-linkage UPGMA tree [14] (`scipy.cluster.hierarchy.linkage`) built directly from the embedding distance matrix. *T*_ref_ is a UPGMA tree derived from TimeTree5 pairwise divergence times, then pruned to the leaf set of the OG under comparison. Lower NRF indicates closer topological agreement.

#### Silhouette score

Computed on the pooled (or PCA-projected) embedding matrix under Euclidean distance, using taxonomic family as the class label. Families with fewer than two species in an OG are excluded. Higher values indicate tighter intra-family clustering relative to inter-family separation [15].

### C. Consensus Reconstruction Across OGs

In addition to per-OG evaluation, we constructed a level-specific consensus tree to summarise the phylogenetic structure shared across all OGs within a taxonomic benchmark. This analysis was performed only after benchmark evaluation.

For each OG, we computed the pairwise distance matrix among the species using the top-performing embedding strategy selected. Then, the OG-specific distance matrices were averaged element-wise to obtain a consensus distance matrix 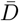 .

A consensus tree was reconstructed from 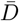 using average-linkage hierarchical clustering (UPGMA) [16]. We evaluated the consensus by computing Mantel *r* against the corresponding TimeTree divergence-time matrix and NRF against the benchmark reference tree, both restricted to the same shared species set.

### D. OGs Saturation Analysis

To determine how many OGs are required for stable reconstruction at each taxonomic scale, we subsample *K* OGs at random, average their pairwise distance matrices over the common species subset, build a UPGMA consensus tree, and compute Mantel *r* and NRF on the resulting tree. The procedure is repeated for *n*_boot_ = 20 bootstrap replicates per value of *K*. The resulting saturation curve defines the minimum OG count needed for reliable distance-based and topological inference.

### E. Evolutionary Rate Analysis

OrthoDB v12 supplies evolutionary-rate estimates per-OG, derived from pairwise *d*_*N*_ */d*_*S*_ ratios, across member species [7]. We compute Spearman rank correlations [17] between these rates and each of the three evaluation metrics to test whether gene conservation modulates Enformer’s phylogenetic signal, and whether any such relationship strengthens at broader taxonomic scales.

### F. Tanglegram

A tanglegram is a visualisation tool used to compare two phylogenetic trees [18]. The two trees are drawn facing each other, with their leaves connected by horizontal lines; the degree of crossing among those lines reflects the topological discordance between the two hypotheses. The reference tree, for this article, was taken from TimeTree5 [9] (i.e. a curated, time-calibrated phylogeny source) and placed against the tree inferred from our Enformer embeddings.

The inspection of the resulting tanglegram provides an intuitive assessment of where the two topologies agree and where they do not, complementing the quantitative measures reported by the Mantel test and the Robinson–Foulds distance with an immediate visual proof.

It was used for Primates and Plants (see Figure 1 and Figure 3) but not for Vertebrates because of the high number of species involved.

**Fig. 1.**
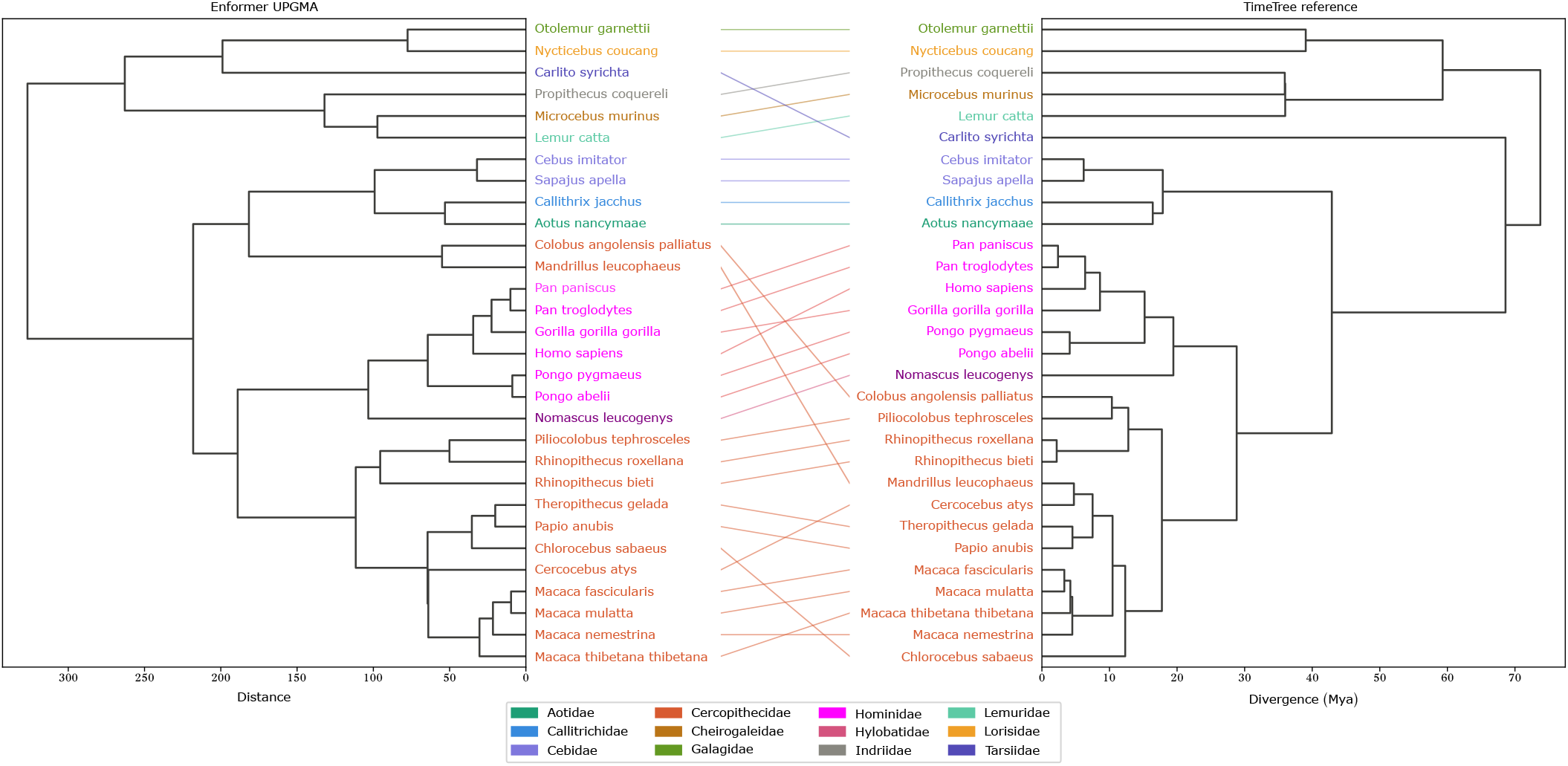
Tanglegram of the Enformer UPGMA consensus tree (left, distance axis) versus the TimeTree5 reference (right, divergence in Mya) for 30 primate species (4 dropped because not present in TimeTree5). Coloured connecting lines link the same species across the two dendrograms; crossing lines indicate topological discrepancies. Leaf colours indicate family membership as shown in the legend.

## III. RESULTS

### A. Primate Benchmark and Strategy Optimisation

#### 1) Dataset

The OrthoDB v12 query at the Primate node (taxon 9443), with universal and single-copy filters, returns 2469 OGs spanning 55 species (86,526 gene-level entries). After coordinate retrieval, gene-length filtering (≤114,688 bp), and retention of only universally represented OGs, **702 OGs** across **34 species** remain, yielding 23,868 (OG, species) windows. Strategy selection is conducted on a pilot of 128 OGs; the winning configuration is then re-evaluated on the full 702-OGs corpus in Section III-B. With 34 species, spanning ≤ 74 Mya of divergence, and a dense, well-characterised TimeTree reference, they are the taxonomic scale closest to Enformer’s training distribution, and thus the expected performance ceiling.

#### 2) Flanking Regulatory Context Carries the Signal

Table I compares all pooling strategies, tested on the 128-OGs pilot under Euclidean distance. The difference between strategy A and strategies B and C is the pooling size: A pools across the full 896-bin window, and it progressively shrinks first to 448 bins and then to gene-proximal bins (≈ 10–50 bins).

**TABLE I.** Strategy comparison on 128 Primate OGS (Euclidean distance). Values: mean ± std across OGS. Lower NRF, higher Mantel *r* / silhouette = better.

| Strategy | Mantel $r$ | NRF | Silhouette |
| --- | --- | --- | --- |
| A pca-16 per-OG | **0.739 ± 0.166** | 0.613 ± 0.111 | **0.223 ± 0.084** |
| A pca-32 per-OG | 0.724 ± 0.179 | **0.587 ± 0.108** | 0.126 ± 0.058 |
| A raw (3072) | 0.643 ± 0.215 | 0.616 ± 0.108 | 0.134 ± 0.094 |
| D mean+std | 0.590 ± 0.206 | 0.643 ± 0.103 | 0.126 ± 0.086 |
| A pca-32 global | 0.599 ± 0.214 | 0.705 ± 0.093 | 0.087 ± 0.125 |
| E max all | 0.481 ± 0.193 | 0.692 ± 0.092 | 0.106 ± 0.067 |
| B central 448 | 0.355 ± 0.205 | 0.788 ± 0.078 | $-0.008 \pm 0.090$ |
| C gene bins | 0.301 ± 0.206 | 0.839 ± 0.074 | $-0.020 \pm 0.078$ |

Strategy C (gene-overlapping bins only) is the weakest configuration tested, at Mantel *r* = 0.301± 0.206 and silhouette − 0.020 ± 0.078. Strategy B (central 448 bins) follows C with a small improvement: it achieves Mantel *r* = 0.355±0.205 with a marginally negative silhouette (−0.008±0.090). However, both of them fall far below strategy A without PCA (Mantel *r* = 0.643±0.215), and far below the best strategy A variant (per-OG PCA-16, *r* = 0.739 ± 0.166).

#### 3) Per-OG PCA Outperforms Global Projection

Within strategy A, per-OG PCA consistently outperforms both the raw 3072-dimensional embeddings and the global-PCA alternative (Table I). Sixteen components achieve the best Mantel *r* (0.739± 0.166) and silhouette (0.223 ± 0.084); 32 components achieve the best NRF (0.587 ± 0.108). Per-OG variance analysis confirms that 90% of explained variance is captured within ≈ 17 components (median across OGs), consistent with 16 being near-optimal.

#### 4) Distance Metric Has Modest Effect

Table II reports the three distance metrics under strategy A raw and PCA-16. Under both strategies, Euclidean displays the highest result on Mantel *r* (0.643 and 0.739), while all three distances converge on close NRF. Silhouette is not present in the table since it is independent of the distance metric. Distance metric is shown to be the least impactful hyperparameter in the pipeline, especially under raw: pooling and PCA account for far greater variance in the outcome. Euclidean is adopted as the default for all subsequent analyses.

**TABLE II.** Distance metric comparison for strategy A (128 Primate OGS), using raw features and pca-16 representations.

| Representation | Metric | Mantel $r$ | NRF |
| --- | --- | --- | --- |
| Raw | Euclidean | **0.643 ± 0.215** | 0.616 ± 0.108 |
| | Correlation | 0.629 ± 0.234 | **0.604 ± 0.112** |
| | Cosine | 0.621 ± 0.233 | 0.606 ± 0.112 |
| PCA-16 | Euclidean | **0.739 ± 0.166** | 0.613 ± 0.111 |
| | Correlation | 0.588 ± 0.123 | 0.560 ± 0.118 |
| | Cosine | 0.593 ± 0.125 | **0.555 ± 0.119** |

#### 5) Number of OGs vs. Reconstruction Quality

Table III reports saturation curves on the 128-OGs pilot. Mantel *r* rises steeply from 0.832 at *K* = 5 and plateaus by *K* = 50; the gain from 50 to 128 OGs is only 0.004. NRF also continues to improve more gradually. Variance, instead, collapses rapidly: the standard deviation *σ* of Mantel *r* falls from 0.050 at *K* = 5 to 0.006 at *K* = 100.

**TABLE III.** K-OGS saturation (strategy A, per-og pca-16, Euclidean, 20 bootstrap replicates per *K*).

| $K$ | Mantel $r$ | $\sigma$ | NRF | $\sigma$ |
| --- | --- | --- | --- | --- |
| 5 | 0.832 | 0.050 | 0.535 | 0.086 |
| 10 | 0.856 | 0.026 | 0.506 | 0.058 |
| 20 | 0.877 | 0.021 | 0.467 | 0.035 |
| 50 | 0.881 | 0.016 | 0.454 | 0.034 |
| 100 | 0.884 | 0.006 | 0.459 | 0.019 |
| 128 | 0.885 | — | 0.444 | — |

**TABLE IV.** Primates reconstruction quality (702 ogs, 34 species)

| Metric | Per-OG mean | Consensus |
| --- | --- | --- |
| Mantel $r$ | $0.757 \pm 0.139$ | **0.902** |
| NRF | $0.577 \pm 0.096$ | 0.481 |

**TABLE V.** Vertebrate reconstruction quality (83 ogs, 150 species)

| Metric | Per-OG mean | Consensus |
| --- | --- | --- |
| Mantel $r$ | $0.618 \pm 0.056$ | **0.752** |
| NRF | $0.749 \pm 0.031$ | 0.606 |

**TABLE VI.** Plant reconstruction quality (92 ogs, 40 species)

| Metric | Per-OG mean | Consensus |
| --- | --- | --- |
| Mantel $r$ | $0.722 \pm 0.042$ | **0.803** |
| NRF | $0.830 \pm 0.049$ | 0.647 |
| Silhouette | 0.158 | — |

Therefore, *K* ≈ 50 OGs is preferable for stable distance-based analysis, while *K* ≥ 100 for topological accuracy. These thresholds inform OG sampling at broader taxonomic levels, where the number of universal single-copy OGs can be considerably smaller.

Based on this sweep, we adopt the following configuration for all subsequent taxonomic levels: **strategy A, per-OG PCA-16, Euclidean distance**.

### B. Full Primate Dataset

The consensus reconstruction substantially improved performance relative to individual OG reconstructions. Whereas per-OG analyses yielded a mean Mantel correlation of 0.757 ± 0.139, the consensus tree reached 0.902, indicating that averaging signal across loci recovered a species-distance structure much more consistent with the reference divergence pattern. Consensus also reduced topological error, with NRF decreasing from a per-OG mean of 0.577 ± 0.096 to 0.481. Together, these results indicate that although individual OGs contain variable and sometimes noisy phylogenetic signal, aggregating information across many OGs enhances recovery of the shared primate phylogenetic structure.

#### 1) Consensus tree vs TimeTree5

Figure 1 shows a tanglegram comparing the UPGMA consensus tree (averaged over all 702 OGs) with the TimeTree5 reference. The major clades (Hominidae, Cercopithecidae, Platyrrhini, and the strepsirrhine-tarsier outgroup) are recovered correctly. The most visible discrepancies are the placements of *Colobus angolensis palliatus, Mandrillus leucophaeus*, and *Chlorocebus sabaeus*. Residual crossings within Cercopithecidae are consistent with the consensus NRF of 0.481: higher-level clade structure is resolved correctly, with uncertainty concentrated at short internodes where divergence times are compressed.

#### 2) n-OGs saturation: full corpus

Figure 2 extends the saturation analysis to the full 702-OGs range.

**Fig. 2.**
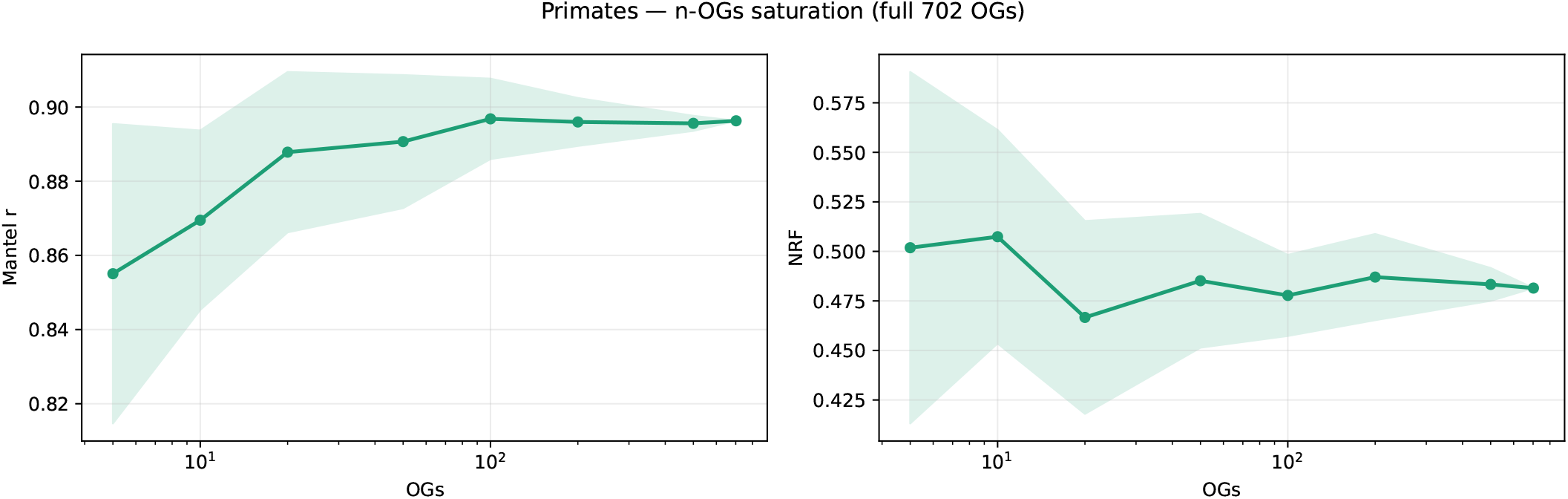
n-OGs saturation curve for Primates (702 OGs, 34 species). Each point is the mean over 20 bootstrap replicates; the shaded band shows ± 1 *σ*. Left: Mantel *r*; right: NRF. Both axes use a log scale on the OG count.

**Fig. 3.**
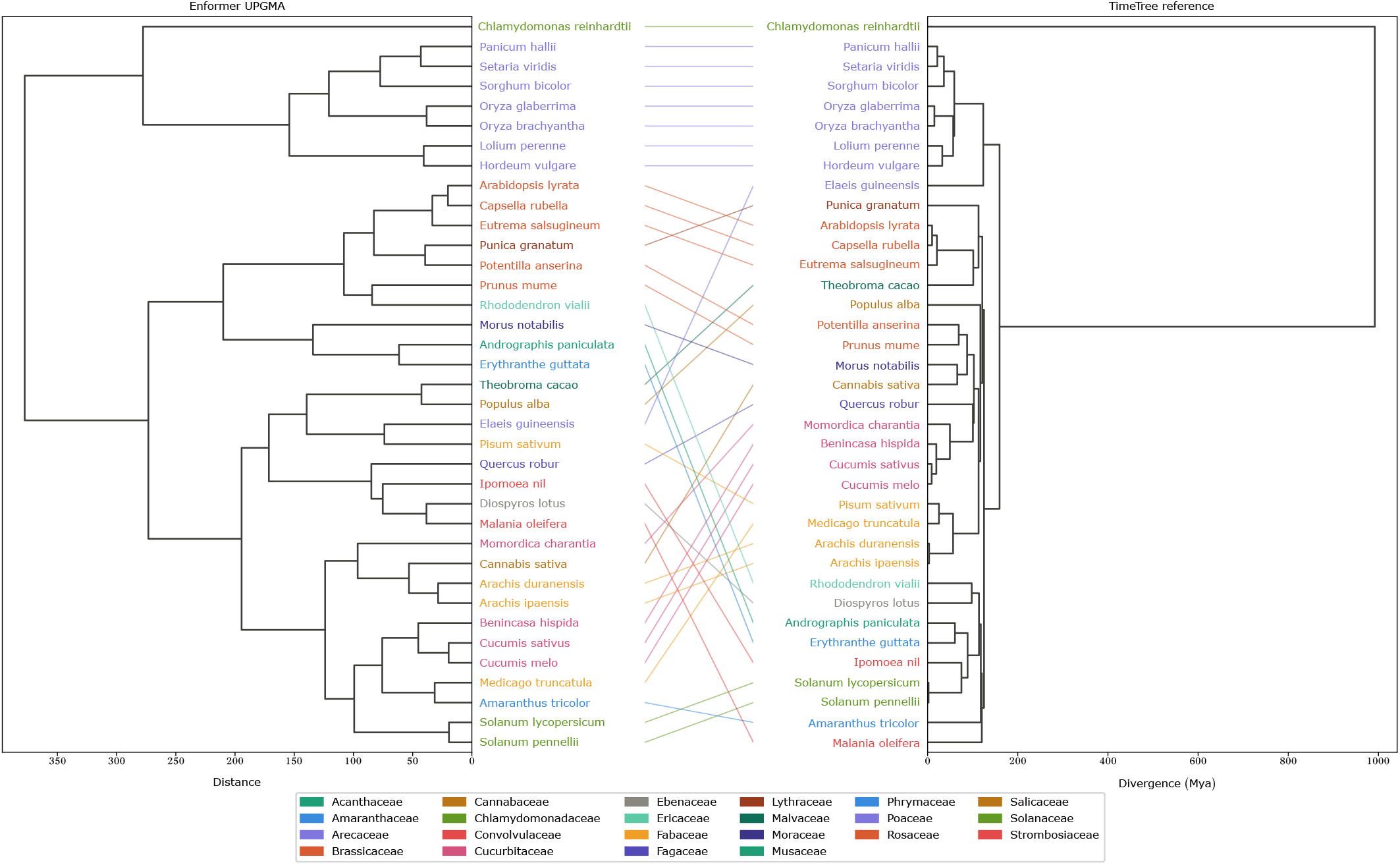
Tanglegram of the Enformer UPGMA consensus tree (left, distance axis) versus the TimeTree5 reference (right, divergence in Mya) for 37 plant species (3 dropped because not present in TimeTree5). Coloured connecting lines link the same species across the two dendrograms; crossing lines indicate topological discrepancies. Leaf colours indicate family membership as shown in the legend.

Consistent with the results already shown above, Mantel *r* plateaus by *K* ≈ 30 (*r* ≈ 0.889) and changes by less than 0.005 between *K* = 50 and *K* = 702. NRF decreases steadily from 0.500 at *K* = 5 to≈ 0.482 at *K* = 702, with variance collapsing as the OG count grows.

This confirms the pilot-scale conclusion: *K* ≈ 50 suffices for distance-based analysis, and *K* ≥ 100 is preferable for topological accuracy, a threshold comfortably exceeded by the 702-OGs primate corpus.

#### 3) Evolutionary rate vs. signal quality

On the 128-OGs pilot, OrthoDB evolutionary rate was weakly negatively correlated with Mantel *r* (*ρ* =− 0.20, *p* = 0.026), suggesting that slowly-evolving genes produce marginally better embeddings. This effect is lost with scaling-up: on the full 702-OGs corpus, *ρ* = 0.005 (*p* = 0.897) for Mantel *r* and *ρ* = − 0.045 (*p* = 0.231) for NRF. Both Mantel *r* and NRF are consistent with zero, showing no sign of correlation between the evolutionary rate and the performance metrics. Within the primate clade, where evolutionary rates cluster near 1.0, and sequence divergence is modest, embedding quality is therefore robust to variation in gene conservation rate. Therefore, the 128-OGs trend is best interpreted as a sampling artefact.

### C. Vertebrate Generalisation

Querying OrthoDB v12 at the Vertebrata node (taxon 7742) with universal and single-copy filters, followed by coordinate retrieval and gene-length filtering, yields **83 OGs** across **150 species**, spanning up to ≤ 450 Mya of divergence time.

Vertebrate embeddings show intermediate performance: consensus Mantel *r* drops to 0.752, from the Primate ceiling of 0.902. The result reflects a reduced regulatory conservation across the deeper evolutionary distances within Vertebrata (spanning all the way to ray-finned fish, ∼ 450 Mya from mammals). Topological accuracy increases compared to the Primate standard (Vertebrate NRF = 0.606 vs Primate NRF = 0.481); this is also consistent with the increase of out-of-distribution operation, as the nucleotide sequences diverge further from human and mouse.

The saturation analysis is consistent with the Primate results: the vertebrate dataset reaches stability in Mantel *r* and NRF by approximately *K* ≈ 50 OGs. The 83-OGs vertebrate set comfortably exceeds this threshold.

Also for Vertebrate group species, the OrthoDB evolutionary rate shows no correlation with Mantel *r* (*ρ* = − 0.096, *p* = 0.386, *n* = 83) or NRF (*ρ* = 0.068, *p* = 0.544, *n* = 83); a further confirmation that gene conservation rate does not systematically predict embedding quality.

### D. Lateral Generalisation — Plants (Viridiplantae)

The Plants macrogroup represents the most challenging test for our pipeline, for a structural reason: plant genomes lack the chromatin architecture and distal-enhancer repertoire that Enformer was trained to interpret. Given that Viridiplantae diverged from the animal lineage approximately 1,500 Mya, a near-complete collapse of phylogenetic signal is therefore expected *a priori*.

Querying OrthoDB at the Viridiplantae node (taxon 33090) with the same universal and single-copy filters, followed by coordinate retrieval and gene-length filtering, yields **92 OGs** across **40 species**. The Primate-optimal configuration (strategy A, per-OG PCA-16, Euclidean) is applied unchanged.

Contrary to our null expectation, the plant consensus embeddings do retain a substantial distance-to-divergence correlation (Mantel *r* = 0.803), only modestly below the Primate ceiling (0.902). Topological accuracy, however, degrades sharply: NRF rises from 0.481 in Primates to 0.647 in plants. The Spearman correlation between OrthoDB evolutionary rate and Mantel *r* is *ρ* = 0.103 (*p* = 0.330, *n* = 92); the correlation with NRF is *ρ* = 0.020 (*p* = 0.848).

## VI. DISCUSSION

### A. The Taxonomic Radius of Enformer

Table VII summarises the above results across all the taxonomic levels under analysis.

**TABLE VII.** Cross-scale reconstruction quality summary.

| Taxon | Div. (Mya) | Mantel $r$ | NRF | Silhouette |
| --- | --- | --- | --- | --- |
| Primates | $\leq 74$ | 0.902 | 0.481 | 0.133 |
| Vertebrates | $\leq 450$ | 0.752 | 0.606 | 0.181* |
| Plants | $\leq 1500$ | 0.803 | 0.647 | 0.158 |

All results use strategy A, per-OG PCA-16, Euclidean. Values are consensus scores over the full OGs corpus at each level. Silhouette is reported as the per-OG mean.
\* silhouette is computed using families for Primates and Plants, but using orders for Vertebrates, since it is a higher-level grouping.

Mantel *r* follows a non-monotonic pattern across scales (0.902 → 0.752 → 0.803), declining from Primates to Vertebrates before partially recovering at the Plant level, and warrants careful interpretation. We identify three plausible, non-mutually-exclusive contributors:

#### (i) divergence-structure effect

Despite the nominal ≤ 1500 Mya ceiling, the plant species distribution is heavily compressed, with the exception of *Chlamydomonas reinhardtii*, which diverged ∼ 1000 Mya; the remaining 39 species cluster within ≤ 200 Mya of one another. The effective divergence range of most plant pairs is therefore considerably narrower than the one in the Vertebrate dataset, where species span the full ≤ 450 Mya window relatively uniformly. Pairwise distance ordering within a tight divergence band is an intrinsically easier task for the Mantel test, which may explain the recovered *r* despite worse topology.

#### (ii) dataset size effect

With only 40 species compared to 150 vertebrates, the plant distance matrix contains fewer closely-spaced internodes, reducing opportunities for local misordering to depress Mantel *r*.

#### (iv) OGs selection effect

The 92 plant OGs retrieved under the universal single-copy filter may be enriched for deeply conserved housekeeping loci whose surrounding sequence context retains coarser-grained compositional structure detectable by Enformer even far out of distribution.

Instead, the NRF gradient (0.481 → 0.606 → 0.647) is strictly increasing, meaning that topological accuracy degrades consistently with evolutionary distance from the training distribution. This could be explained by the fact that Vertebrates, being phylogenetically closer to Enformer’s two training genomes (human and mouse), retain more of the chromatin architecture and regulatory grammar that the model was explicitly trained to interpret, thus yielding better topological resolution than Plants despite the latter’s narrower effective divergence range.

The dissociation between Mantel *r* and NRF reflects two genuinely distinct aspects of reconstruction quality. Mantel *r* is dominated by large-scale distance structure: well-separated species pairs contribute disproportionately to the rank correlation, and it is precisely these coarse, deep divergences that become unreliable far out of distribution. NRF, by contrast, depends on the correctness of individual bipartitions at all depths; closely related species are still placed near one another in embedding space, but the broader branching order above them degrades, a pattern clearly visible in the Plant tanglegram. Fine-grained local clustering therefore persists longest, while coarse topological structure fails first.

### B. Regulatory Context as a Phylogenetic Signal

A central finding concerns the localisation of phylogenetic signal within Enformer embeddings. Restricting the pooling to the gene locus alone (strategy C) causes the Mantel *r* to collapse from 0.739 to 0.301 on the 128-OGs Primate pilot: a drop that pinpoints the flanking regulatory context, rather than the coding sequence itself, as the primary carrier of phylogenetic information.

A natural confound is that strategy C averages over far fewer bins (≈ 10 − 50, gene-length dependent) than strategy A (896 bins), so the collapse in Mantel *r* could reflect increased estimator variance, rather than a genuine loss of biological signal.

However, two observations argue against this interpretation. Strategy B, which pools the central 448 bins, achieves only a modest recovery (*r* = 0.355), well below the optimal strategy A (*r* = 0.643). Indeed, if bin count were the primary driver, a smoother gradient across A, B, and C would be expected. Moreover, the silhouette score under strategy C is negative (− 0.020 ± 0.078), indicating that family-level clustering is actively worse than random, a pattern inconsistent with a mere variance inflation and instead suggestive of a qualitative absence of discriminative signal. Together, these observations support the interpretation that the drop reflects the removal of regulatory content rather than a statistical artefact.

This result is not unexpected. Enformer was explicitly trained to interpret long-range regulatory interactions, chromatin topology, and enhancer repertoires; excising the surrounding genomic context removes precisely the features the model was designed to encode.

Unlike classical marker-gene approaches, Enformer-based phylogenetics compares the regulatory landscape *surrounding* those sequences as learnt by a deep neural network. The two signals are complementary: sequence phylogenetics reads synonymous and non-synonymous substitution rates in coding regions, whereas regulatory phylogenetics captures the evolution of non-coding regulatory elements at scale. Integrating both signals within a unified tree-inference framework is a natural direction for future work.

An alternative, more data-centric interpretation leads to a similar conclusion by a different route. The genes embedded in this study are universal single-copy orthologs, conserved by construction across all species in the dataset. Their coding sequences and core promoter elements are expected to be highly similar across taxa, carrying little species-discriminating signal by design. Strategy C, by restricting pooling to the gene-overlapping bins, reads precisely the most conserved part of the window, which is the region where sequences look most alike because of the orthology constraint itself. The flanking context, by contrast, evolves more freely: enhancers, insulators, and chromatin domain boundaries accumulate lineage-specific variation under weaker purifying selection, and it is this faster-evolving periphery that strategy A captures.

This interpretation is independent of Enformer’s architecture and training objectives, yet arrives at the same outcome: gene-overlapping bins are uninformative not only because Enformer was not trained to read them in isolation, but because the sequences themselves are too similar across species to carry a divergence signal.

### C. Per-OG PCA and the Gene-Specific Latent Space

Another result is the advantage of per-OG PCA over global PCA: it reveals that each orthologous gene occupies a distinct principal-variance structure in Enformer space. Global PCA imposes a single shared basis across all 23, 868 rows, averaging over gene-specific effects and degrading the within-OG distance geometry, on which phylogenetic inference depends. Per-OG PCA preserves that local structure, raising silhouette from 0.087 (global) to 0.126, improving Mantel *r* from 0.599 to 0.724 and NRF from 0.705 to 0.587. The improvement from 3072 to 16 dimensions with PCA is consistent with per-OG variance analysis showing that ∼ 90% of explained variance is captured within ≈ 17 components (median across OGs), confirming that 16 is near-optimal and that the gain reflects genuine dimensionality reduction rather than overfitting to a small component count.

This finding has a practical implication for cross-scale studies. Universal single-copy OG counts shrink rapidly as taxonomic queries broaden, but per-OG PCA remains well-defined even with small OGs sets, because each fit uses only the ∼ 30 − 150 species within an individual OG. The winning configuration, therefore, transfers cleanly to regimes where global-PCA approaches would be starved of data.

The consensus reconstruction should be interpreted as a summary of shared multigene structure rather than a formal phylogenetic estimate. Averaging OG-specific distance matrices before UPGMA reconstruction captures species relationships that are repeatedly supported across loci while reducing OG-specific noise; pairwise distances are also more comparable across independently fitted PCA spaces than the component axes themselves.

However, three limitations apply: OG-specific distance matrices may differ in scale, potentially giving disproportionate weight to loci with greater dispersion; restricting to species shared across all OGs reduces taxon coverage; and averaging across loci may smooth over genuine gene-tree discordance.

### D. Evolutionary Rate

A natural hypothesis entering this analysis was that slowly evolving genes are more informative embeddings and would provide better phylogenetic reconstruction. Under this view, a positive correlation between OrthoDB evolutionary rate and NRF (or a negative one with Mantel *r*) would be expected, strengthening as the model moves further out of distribution and sequence divergence accumulates.

However, the data do not support this hypothesis. On the 128-OGs Primate pilot, a weak negative association with Mantel *r* is observed (*ρ* = − 0.20, *p* = 0.026), which could superficially be read as confirming the prediction. However, this effect disappears entirely on the full 702-OGs corpus (*ρ* = 0.005, *p* = 0.897), strongly suggesting it was a sampling artefact of the pilot: with only 128 OGs drawn from a pool of 702, the subset happened to contain a slight enrichment of slowly-evolving genes with above-average embedding quality, a fluctuation that vanishes at scale. The hypothesis also fails at broader taxonomic scales, where rate variance is larger, and the model is more out-of-distribution; precisely the regime where the effect should be strongest, if conservation drove embedding quality.

This implies that what makes a gene informative for Enformer-based phylogenetics is not its rate of sequence evolution per se, but more likely properties of its surrounding regulatory landscape, such as the density, diversity, and cross-species conservation of its flanking enhancer repertoire, which are not captured by *d*_*N*_ */d*_*S*_ alone and remain an open question for future investigation.

### E. Limitations and Future Directions

Before detailing the limitations of this work, it is important to clarify once again its scope. This paper does not aim to establish a new benchmark nor to propose Enformer as a competitive phylogenetic inference tool; rather, it assesses whether Enformer can learn representations that reflect evolutionary relationships across the broader tree of life, even though it was trained only on human and mouse genomes. The experiments are therefore designed as a proof-of-concept of Enformer’s generalisation capacity, not as an optimisation of a phylogenetic pipeline. The absolute performance values reported here should be interpreted in this light: they characterise the boundary of what an unmodified, out-of-the-box regulatory model can see across evolutionary time, rather than what a purpose-built phylogenetic tool could achieve.

#### Training distribution

Enformer was trained exclusively on human and mouse genomes, meaning that every cross-species inference in this work is out-of-distribution to some degree, ranging from the minimal shift of the great apes to the near-complete breakdown expected for organisms without chromatin-based regulation. While our results demonstrate that useful phylogenetic signal persists well beyond the training distribution, the absolute performance ceiling is set by this constraint. Fine-tuning on a phylogenetically diverse set of reference genomes, spanning multiple vertebrate classes or plant lineages, would directly extend the model’s regulatory vocabulary and test how much of the observed signal degradation is attributable to the training distribution rather than to the architecture itself, though this would be computationally expensive.

#### Orthology constraint

The universal single-copy filter, while necessary to ensure cross-species comparability, imposes a strong conservation bias on the gene set. Relaxing the orthology constraint to allow partially populated OGs, where some species are missing, would substantially increase the available gene set, though this would require imputation or missing-data-aware tree reconstruction methods to avoid introducing systematic bias in the placement of underrepresented clades. Future gains in reconstruction quality may therefore come less from brute-force addition of OGs beyond *K* ≈ 100 and more from qualitative improvements to the gene panel, for instance, through targeted curation selecting loci embedded in informative regulatory landscapes.

#### UPGMA

All trees in this work are inferred by UPGMA, which assumes equal substitution rates across all lineages. This assumption is well-supported within Primates, where rate variation is modest, but becomes increasingly questionable at the Vertebrate and Plant scales. Replacing UPGMA with a rate-heterogeneity-aware method could recover topological accuracy currently lost to this assumption; comparing both methods on the same consensus distance matrices would directly quantify this contribution without requiring any additional model runs.

#### Mechanistic interpretation and signal attribution

While this work demonstrates that flanking regulatory context rather than the gene body carries the phylogenetic signal, as shown by the contrast between strategies A and C, it does not explain which specific features drive this effect. It remains unclear whether the model has implicitly learnt a representation of regulatory conservation or whether the signal emerges from more indirect properties of sequence composition that happen to correlate with divergence time. Attribution methods such as in-silico mutagenesis or attention-weight analysis could identify which specific regulatory elements (enhancers, CTCF binding sites, or broader chromatin architecture) are the primary drivers, and whether the effect is uniform across OGs or concentrated in gene classes embedded in particularly informative regulatory landscapes.

## Acknowledgment

N.S. is supported by the Wellcome Trust grant (225220/Z/22/Z). S.G.R. is supported by the MRC grant (MC UU 00029/3). J.R.H. is supported by the Wellcome Trust grants (225220/Z/22/Z and 106130/Z/14/Z) and the MRC grant (MC UU 00029/3).

## Declaration

J.R.H. is a co-founder and director of Nucleome Therapeutics and provides consultancy to the company.

